# Characterizing the Effects of Protein Glycosylation Perturbation on Phosphorylation Signaling

**DOI:** 10.64898/2025.12.18.695253

**Authors:** Effram Wei, Hongyi Liu, Michael Betenbaugh, Hui Zhang

## Abstract

Protein glycosylation and phosphorylation constitute two pervasive regulatory layers in mammalian cells, yet the effects that protein glycosylation play in phosphorylation signaling remain poorly understood. Here we show that controlled perturbation of N-linked glycan biosynthesis through glycoengineering fundamentally rewires phosphorylation signaling networks in human cells. Using comprehensive proteomics approaches, we simultaneously profiled the global proteome, glycoproteome, and phosphoproteome in engineered HEK293 cells designed to eliminate core fucosylation while enhancing sialylation and reducing GlcNAc branching complexity. Glycoengineering emerged as the dominant source of molecular variation across all datasets, with over 9,800 intact glycopeptides identified of which 3,400 are significantly altered, establishing a remodeled baseline cellular state. Upon serum stimulation, engineered cells not only exhibited markedly decreased phosphorylation responses compared to wild-type cells, but comprehensively re-wired to prefer signaling away from canonical EGFR/mTOR growth pathways. These findings establish a systematic framework for targeting glycosylation-phosphorylation regulation and nominate glycan-dependent signaling nodes as potential therapeutic vulnerabilities in glycosylation-remodeled disease states.

## INTRODUCTION

Protein glycosylation is a pervasive, enzyme-directed protein modification that decorates the majority of secreted and membrane proteins and regulates folding, trafficking, stability, receptor signaling, and immune recognition in mammalian systems^1,2,3^. Among glycan classes, N-linked glycosylation (covalent attachment of glycans to asparagine in the consensus sequon) expresses on the cell surface or secreted from cells, constitute one of the most well-studied glycosylation forms^4,5^. As arguably the most diverse and abundant protein modifications, glycosylation’s systems-level importance is its clinical significance^6,7,8,9^. In congenital disorders of glycosylation, even modest perturbations in glycan biosynthesis, primarily by glycosyltransferases, can have profound physiological consequences^3,10^. In cancer, tumor cells commonly remodel N-glycan branching, sialylation, and fucosylation, reprogramming receptor–ligand interactions, cell–matrix adhesion, immune surveillance, and growth factor signaling; these alterations are increasingly recognized as both disease drivers and actionable biomarkers^2,11,12,13^. Therefore, protein glycosylation is crucial for their function and required for functional studies.

Protein phosphorylation as perhaps the most extensively studied protein post-translational modification (PTM) controls almost every aspect of cell signaling and transduction through the attachment and removal of a phosphate group on primarily serine (S), threonine (T), and tyrosine (Y) amino acid residues^14,15,16,17^. The human genome comprises of over 500 highly specific protein kinases that, together with approximately 200 protein phosphatases, and with diverse substrates and phosphorylation sites, orchestrate dynamic and reversible changes in protein activity, localization, and interaction networks across pathways governing proliferation, metabolism, and stress responses^17,18^. Dysregulation of phosphorylation mechanisms is frequently implicated in diseases such as cancer, where aberrant kinase activity or mutations in phosphorylation sites contribute to oncogenesis by disrupting normal signal transduction and cellular homeostasis^19^. Targeting dysregulated phosphoproteins has emerged as a promising therapeutic strategy, evidenced by the development and approval of kinase inhibitors drugs^19,20^.

N-linked protein glycosylation and phosphorylation are already two of the most involved regulatory layers in mammalian systems, yet their combined modifications on proteins are responsible for a further increase in proteome complexity. Unlike other protein modifications studied alongside phosphorylation, such as O-GlcNAcylation which predominantly involves S and T residues, N-linked glycosylation lacks direct residue competition with phosphorylation but instead is implicated through signaling effects, including modulation of receptor trafficking, ligand binding affinity, protein stability, and the spatial organization of signaling complexes that regulate phosphorylation-dependent signaling cascades^21,22^. For example, studies have shown that N-glycosylation of programmed death ligand 1 (PD-L1) prevents phosphorylation-dependent proteasomal degradation by glycogen synthase kinase-3 β (GSK3β) and β-transducin repeat-containing proteins (β-TrCP) while promoting its stability and interaction with PD-1 to enable tumor immune evasion, with epidermal growth factor (EGF) signaling further stabilizing PD-L1 through GSK3β inactivation^23,24,25^. Other studies show that G protein-coupled receptors (GPCRs) demonstrate diverse functions of glycosylation on phosphorylation, where N-linked glycosylation in extracellular domains influence phosphorylation-dependent signal bias at intracellular loops, contributing to receptor internalization, functional selectivity, and ligand responsiveness^26,27^. Extensive literature links the aberrant crosstalk between N-glycosylation and itself, as well as site-specific phosphorylation to oncogenic signaling: tumor-associated hyper-branching, terminal sialylation or core fucosylation of N-linked glycans can mask phosphodegrons, disrupt kinase- or β-TrCP recognition, and simultaneously engage lectin/siglec pathways, collectively fueling malignant transformation, metastasis and inflammation^28,29,30^. However, most studies investigating N-glycosylation and phosphorylation rely on biochemical approaches such as PNGase F treatment followed by western blotting, lectin binding assays, or site-directed mutagenesis, which lack the resolution to comprehensively map site-specific modifications across the proteome^23,24,25,26^.

As accompanied by the advancement in protein analytical chemistry and liquid chromatography mass spectrometry (LC-MS) based technologies, proteomics toolboxes can give us a better and more holistic profile of glycosylation and the interactions with phosphorylation^31,32,33^. While robust analytical platforms by quantitative phosphoproteomics has been around for over a decade^34,35,36,37^, recent advancement of site-specific N-glycoproteome profiling has allowed quantitative, site-resolved interrogation of thousands of N-glycosylation sites in a single experiment, thereby opening the possibility of monitoring glycan remodeling events in parallel with other post-translational modifications in an unbiased manner^33,38,39,40,41,42,43,44,45,46,47,48,49,50,51^. However, since N-glycosylation is widespread among glycoproteins, any modification in N-glycan biosynthesis can influence numerous cellular pathways. This broad impact makes it challenging to relate physiological changes to altered glycosylation of particular protein groups^52,53^. Additionally, the current research in functions of N-glycosylation on phosphorylation lacks systematic, causal maps that connect defined glycosylation perturbations to network-wide phosphorylation dynamics and kinase activity states in a controlled cellular context^26,33,54,55,56^.

In this study, we employ glyco-gene engineering approaches as a controlled source of intrinsic perturbation on the function of various glycosyltransferases, assessing the impacts of glycosylation engineering on the subsequent N-glycoproteome in human embryonic kidney (HEK293) cells^57,58^. We selected HEK293 cells rather than a clinically derived cell line due to its comprehensive human glycosylation machinery, strong CRISPR knockout efficiency, and deep proteome coverage^59,60,61^. In addition, HEK293 cells do not contain the confounding mutations and heterogeneity in cancer models, while capable of informing the same clinically relevant glycosyltransferase repertoire and phosphorylation signaling pathways, making it suitable for a perturbation study^62,63^.

Particular up or down regulated glycan features in our engineered cell line include fucose, sialic acid, and high N-acetylglucosamine antenna branching, which have been extensively shown to correlate with various diseases clinically, including cancer^64,65,66,67,68,69^. We then subjected these engineered cells, along with wild-type (WT) HEK293 cells, to serum starvation and re-stimulation *in vitro*, which is known to reveal the dynamic changes in phosphorylation activity^70,71,72^. Through integrated proteomics on these cell lines, we were able to monitor the changes in protein expression, the glycoproteome, and the phosphoproteome for both cell lines, as well as their differences under stimulatory conditions. This approach gives a constructive portrayal of primary changes including protein expression of these glycoengineered enzymes and glycan modifications due to genetic perturbation, as well as the secondary changes including the transient response framework in phosphorylation signaling under external stimuli. In doing so, we test the hypothesis that remodeling N-linked glycosylation rewires cellular signaling networks in ways that expose therapeutic vulnerabilities, including those in phosphorylation pathways that lack directly druggable nodes.

To our knowledge, this is the first study to investigate the functions of glycosylation on phosphorylation signaling using glycoengineering of human cells by glycosylation enzyme alterations. Our data showed the changes in glycan biosynthesis propagate through kinase-mediated signaling to reshape pathway activity. By integrating glycosylation and phosphorylation readouts under tightly controlled perturbations, our study establishes a standardized framework for delineating glycosylation–phosphorylation regulation at scale and for nominating glycosylation enzymes, glycan epitopes, or signaling nodes as candidates for therapeutic intervention and target development in glycosylation-remodeled disease states.

## RESULTS

### Proteome-wide effects of glycoengineering

A glycoengineering, serum stimulation study, and integrated proteomics workflow was employed to investigate the effect of glycoengineering on proteins and glycoengineered enzyme expression, as well as the consequent glycosylation changes and their impacts on phosphorylation signaling (**Fig. 1A**). We utilized genetic engineering methods to alter the biosynthesis of N-glycans. Using CRISPR–Cas9, we re-routed N-glycan processing by engineering numerous glycan biosynthesis related genes (see **Supplementary Fig. 1**), beginning with disabling five endogenous genes: fucosyltransferase 8 (FUT8), GDP-mannose 4,6-dehydratase (GMDS), α-1,3-mannosyl-glycoprotein 4-β-N-acetylglucosaminyltransferase A and B (MGAT4A and MGAT4B), and α-1,6-mannosylglycoprotein 6-β-N-acetylglucosaminyltransferase (MGAT5). Loss of FUT8 together with ablation of the GDP-fucose–biosynthetic enzyme GMDS decreases core fucosylation and other forms of antennary fucosylation, while removal of MGAT4A/B and MGAT5 blocks the β1,4- and β1,6-branching steps that normally give rise to highly branched tri and tetra-antennary structures^73,74,75,76,77^. To further bias flux toward sialylation modified termini with reduced branching, we stably over-expressed α-1,3-mannosyl-glycoprotein 2-β-N-acetylglucosaminyltransferase (MGAT1) and α-1,6-mannosyl-glycoprotein 2-β-N-acetylglucosaminyltransferase (MGAT2), as well as β-1,4-galactosyltransferase 1 (B4GALT1) and ST6 β-galactoside α-2,6-sialyltransferase 1 (ST6GAL1)^78^. We also induced a point mutation on the gene encoding native glucosamine (UDP-N-acetyl)-2-epimerase/N-acetylmannosamine kinase (GNE), converting it to its sialuria variant for overproduction of sialic acid via failed feedback inhibition^79^. The glycoengineering method used is described in further detail by Aliyu L. et al (2025, under review). These modifications confines the biosynthesis pathway to favoring sialylated form of either a classic bi-antennary complex glycan or a hybrid complex intermediate^80,81,82^. The resulting glycoengineered cells is expected to have increased synthesis for N-glycans that are almost exclusively non-fucosylated bi-antennary or hybrid forms, displaying high levels of terminal galactosylation and α2,6-linked sialylation. Hereafter, we refer to this engineered product as the All-Included (AI) cell line.

**Figure 1.**
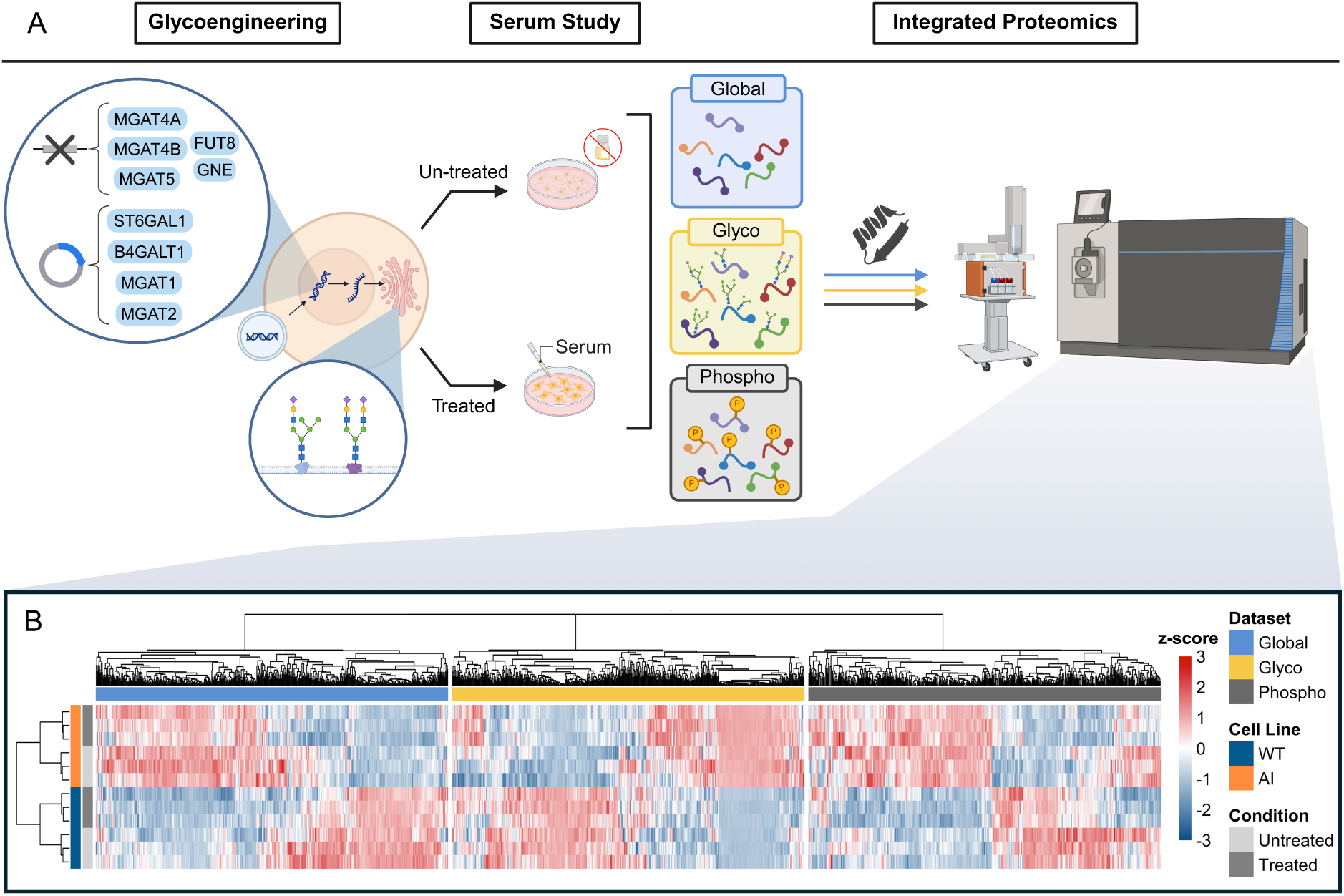
Glycoengineering design and proteome-wide cell line dependent serum stimulation signatures across global proteins, glycopeptides and phosphopeptides. **A.** Schematic of the CRISPR-based glycoengineering strategy and serum-stimulation study followed by integrated global, glyco- and phosphoproteomics. **B.** Cross-dataset differential expression heatmap with WT and AI samples under conditions of starvation (un-treated) and serum-stimulation (treated). The three datasets (in blocks) represent global proteins, intact glycopeptides, and phosphopeptide phosphorylation sites.

After culturing the AI cell line, along with the parental WT HEK293 cell line in complete media, we subjected them to two experimental conditions including starvation in serum free media, and starvation followed by serum re-stimulation (30 minutes window). The samples from each cell line and condition were prepared and analyzed by liquid-chromatography (LC) coupled with data-dependent acquisition (DDA)-tandem mass spectrometry (MS/MS) quantification of the glycoproteome and phosphoproteome. In separate raw expression matrices with duplicates removed, we detected 10924-10927 proteins, 9763-10065 intact-glycopeptides, and 38772-28778 phosphorylation sites (**Supplementary Fig. 2A, Supplementary Table 5**). All three raw datasets were preprocessed with the same workflow: features were cleaned, log2-transformed and PhosR inspired median-centered to mitigate sample-wise intensity shifts, followed by the ensemble imputation method of DreamAI^83,84^. Quality control included coefficient of variation (CV) distribution, Spearman correlation amongst samples, intensity density distribution, and detection threshold by features (**Supplementary Fig. 2B, Supplementary Table 5**).

We generated an integrated view of differential regulation across the global proteome, phosphoproteome and glycoproteome using a cross-dataset heatmap (**Fig. 1B**). We used a Limma based ANOVA framework with 2 × 2 factorial design (cell line × condition), and calculated the moderated F-statistics to identify differentially expressed features that vary across the serum stimulation study (see **Supplementary Table 1**)^85^. For each dataset, features (global proteins, intact-glycopeptides, phosphorylation sites) were ranked by the moderated F-statistic and the top 500 significant features with false discovery rate (FDR) < 0.05 were retained, yielding up to 1,500 features prior to completeness filtering. The corresponding intensity matrix was assembled across all samples, filtered to retain features with at least 70% detected values across samples, and z-score normalized per feature. Samples were hierarchically clustered within each dataset. We observe from the heatmap across all three datasets that samples cluster unsupervised by cell line (rows). A secondary separation is evident specifically in the phosphoproteome, where starvation and serum-stimulated clusters within each cell line. The global proteome shows a more modest serum treatment-driven shift, while the glycoproteome exhibits minimal reorganization over the serum stimulation window.

### Baseline difference from glycoengineered cells under the starved condition

We show the baseline differences due to glycoengineering under the starvation condition, which establishes the causal starting point for our study. Here we restrict the analysis to cells under starvation and compare the AI cell line to the WT cell line. This isolates the effect of glycoengineering on the proteome as well as the glycoproteome, and provides the baseline against which the subsequent serum-induced signaling responses within each cell line should be interpreted. We first analyzed the protein expression of starved cells using DDA-MS results, with the addition of data-independent acquisition (DIA)-MS results, to determine the change in protein expression for glycoengineered cells and confirm successful glycoengineering for the targeted enzymes for N-linked glycan biosynthesis pathway (**Fig. 2A**). The global protein-level readout of the engineered enzymes confirms successful perturbation with all glycoengineered genes being statistically significant (p-value < 0.05) under a one-tailed student’s t-test (**Supplementary Table 2**). Targeted knock-ins (B4GALT1, ST6GAL1, MGAT1, MGAT2) showed elevated protein levels, while knock-outs (GMDS, MGAT4B, MGAT5, FUT8, and MGAT4A) showed significant reductions. The knock-out on FUT8 and MGAT4A was completely successful based on MS detection. The GNE variant, whilst not expected to necessarily change in either direction, shows a modest decrease in protein expression. Notably, our edits induce a loss- or gain-of-function to the target glycoengineered genes.

**Figure 2.**
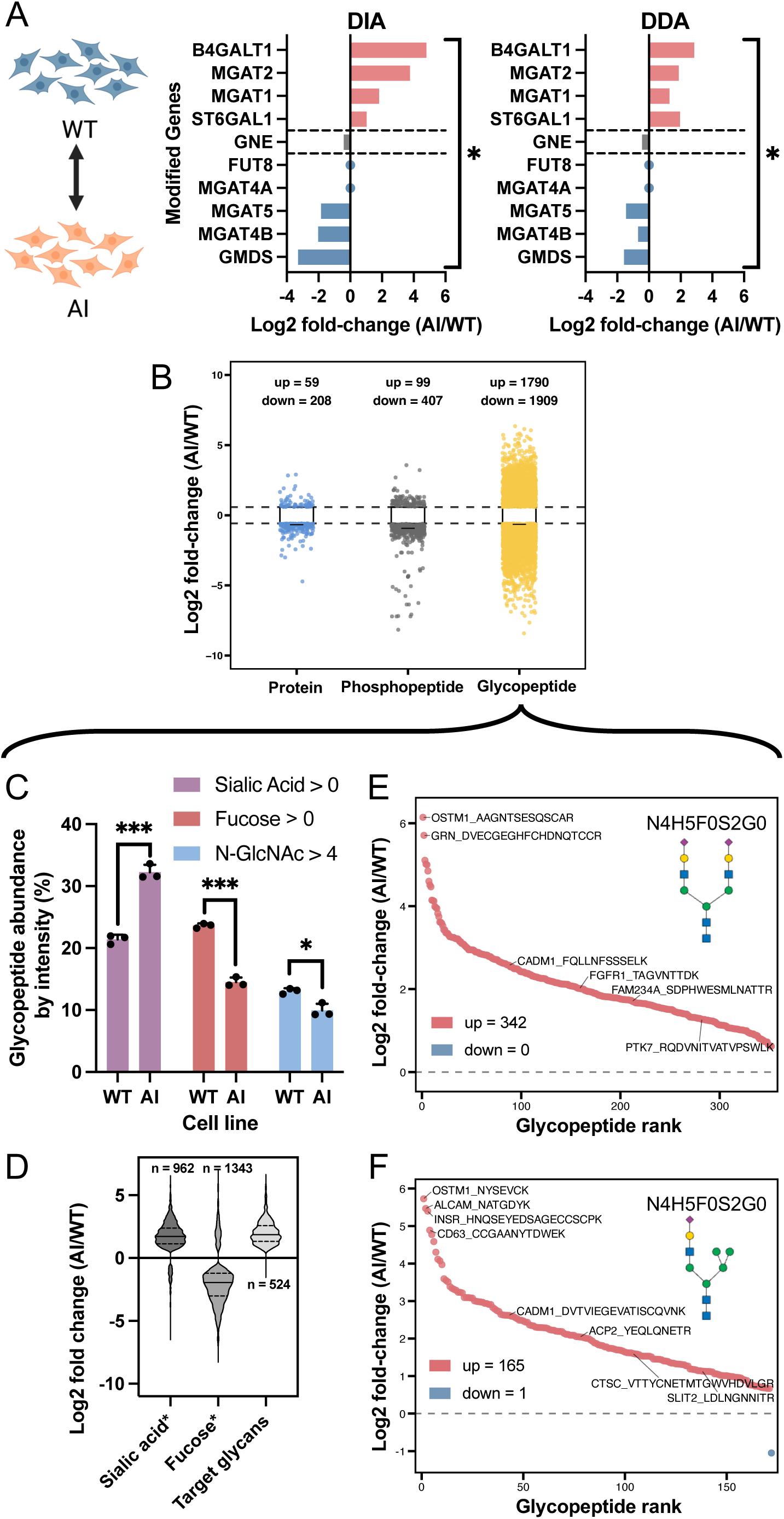
Baseline effect of glycoengineering across proteome layers by comparing AI vs WT cell lines under the starvation condition. **A.** Panels show bar plot of global protein-level changes (indicated by corresponding genes) in target genetically engineered enzymes by both DIA and DDA proteomics. The log2 FC of observed protein regulation was shown in combination with the anticipated genetic engineering of knock-in, knock-out, or mutation. Bracket and * indicates significance of differential expression. **B.** Volcano scatter plot of differential expression analysis on significant global proteins, phosphorylation sites, and intact glycopeptides due to glycoengineering. **C.** Relative abundance of glycopeptides containing specific glycans given by un-transformed ratios, where classes represent glycans containing sialic acid, fucose, and GlcNAc branches (N>4). SEM and significance criteria are shown (*** p-value < 0.001; ** p-value < 0.01; * p-value < 0.05). **D.** Category-wise log2 FC distributions of differentially expressed glycopeptides that contain a sialic acid and no fucose (sialic acid*), contain a fucose and no sialic acid (fucose*), and those that are the target complex/hybrid glycans with absence of high-antennary branching. **E. and F.** S-plots of site-resolved differentially expressed glycopeptides containing the target glycoforms (glycoform notation given by NxHxFxSxGx, where N = GlcNAc, H = hexose, F = fucose, S = Neu5Ac, G = GalNAc), depicting the regulation of **E.** target complex glycan and **F.** target hybrid glycan, both non-fucosylated, α2,6-sialylated, and low antennary branching. Glycopeptides are labelled by detected amino acid sequence and corresponding gene.

We observed that the protein level changes in glycosyltransferase expression were further confirmed by analysis on the N-glycoproteome. To compare the overall effect, we performed pairwise differential expression analysis using an empirical-Bayes linear-model framework in Limma to compare the difference between AI and WT (AI/WT) global proteins, phosphopeptides, and glycopeptides (**Fig. 2B**). Unless otherwise indicated, significance was defined in this study as adjacent p-value (FDR) < 0.05 with |log2 fold-change (FC)| > 0.58 (∼1.5 FC). As expected from a direct manipulation of the glycan biosynthetic machinery, the glycoproteome exhibits by far the largest number of significant baseline feature differences, with additional but more modest changes in the phosphoproteome and global proteome (protein 215; phosphopeptide 339: glycopeptide 3402).

To validate specific changes of glycopeptides at the glycan class level, we quantified relative glycopeptide abundances of the three glycan categories of interest—sialylated (sialic acid > 0), fucosylated (fucose > 0), and branched (N-GlcNAc > 4; tri-, tetra- or higher-antennary)—by un-transformed ratios ratios (total glycan class intensity/total intensity of all glycopeptides) of raw total glycopeptide signal per sample after intensity and detection-rate filtering (**Fig. 2C**). Statistical comparison was made by a two-sample t-test with significance given by *** p-value < 0.001; ** p-value < 0.01; * p-value < 0.05, with SEM and the t-statistic calculated. We saw that AI cells display increased sialylation, decreased fucosylation, and reduced MGAT4/5-dependent high antennary branching, consistent with our glycoengineering design in **Supplementary Fig. S1**. Distributions of log2 FC by glycopeptide type (**Fig. 2D**) show coherent directional shifts (where * species indicates mutually exclusivity): fucosylated species were down-regulated (median ∼ -1.88), whereas sialylated (median ∼ 1.68), and the engineered target glycans according to our schematic (median ∼ 1.85) were up-regulated. We further examine the specific regulation of engineered target glycopeptides in the **Fig. 2E** and **Fig. 2F** s-plots. We annotate representative N-glycoforms using the shorthand notation N_H_F_S_G_, indicating the number of N-acetylglucosamine (GlcNAc, N), hexose residues (mannose/galactose, H), fucose (F), sialic acid (Neu5Ac, S), and N-acetylgalactosamine (GalNAc, G) monosaccharides, respectively. N4H5F0S2G0 represents our engineered target bi-antennary complex glycan with two sialic acids and no fucose, and N3H6F0S1G0 represents the engineered target hybrid glycan with one sialic acid and no fucose. We see that all significant glycopeptides containing our target complex glycan was up-regulated, and only one glycopeptide containing our target hybrid glycan was down-regulated. Furthermore, this up-regulation for both glycoforms appear drastic with a median log2 FC of ∼2. In another s-plot of all effective glycopeptide regulation, we see the lower tail (depleted in AI) is enriched for fucosylated species, while the upper tail (increased in AI) includes the engineered targets or associated targets—non-fucosylated, α2,6-sialylated complex or hybrid forms—demonstrating that genetic edits propagate to individual glycopeptides (**Supplementary Fig. 3**).

The detailed data for the above figures can be found in **Supplementary Table 2**. Conceptually, these baseline results potentially anchor a causal framework such that when cells are subsequently challenged with serum, the divergent phospho-signaling trajectories observed in (**Fig. 1B**) can be interpreted as consequences of this engineered glycan context rather than generic differences between cell lines.

### Glycoengineering remodels the cellular phosphorylation signaling in response to serum stimulation

Serum study datasets underwent the same pre-processing as **Fig. 1B** and was then analyzed using an empirical-Bayes linear model in Limma for differential expression contrasting conditions under serum starvation (un-treated) and serum stimulation (treated) within the AI and WT cell lines separately. The resulting differential expression characterizes the response to serum stimulation by the AI or WT cell line. Statistical significance follows the same criteria and analyses were run independently for the glycoproteome and phosphoproteome. As a first overview, the summary of significant differential phosphorylation sites shows that the serum treatment effect is concentrated in the phosphoproteome, whereas the global proteome and glycoproteome change minimally over the 30 minutes serum stimulation window (**Fig. 3A**). Importantly, the serum response in WT (618 differential features total; 378 up, 240 down) was much greater than that of AI (358 differential features total; 269 up; 89 down). This cell line dependent response to serum stimulation in the phosphoproteome, in combination with **Fig. 2**, may suggest a causal sequence for engineered glycans to the subsequent kinase-mediated signaling responses upon that background.

**Figure 3.**
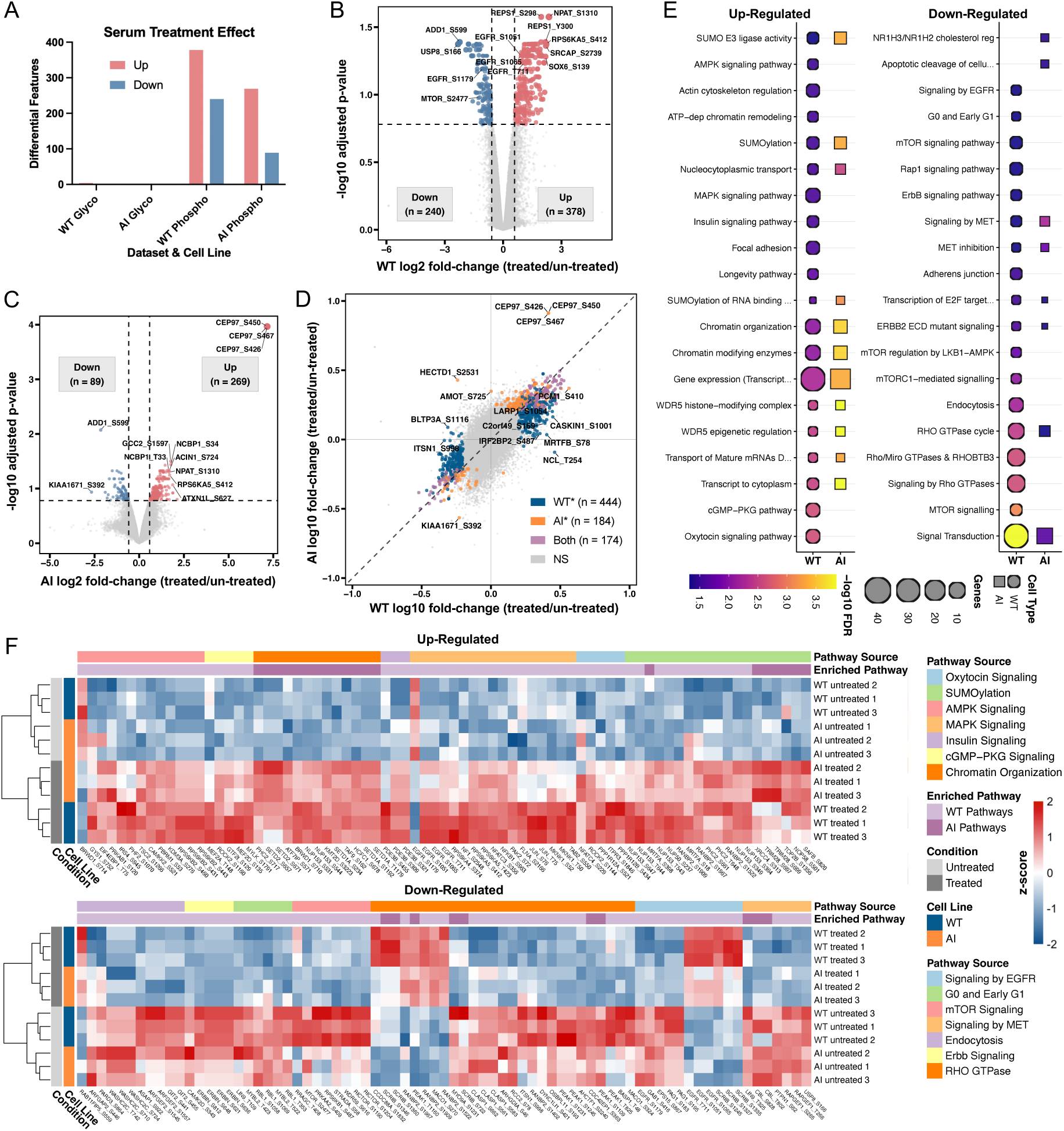
Glycoengineering reprograms serum-induced signaling. **A.** Bar summary of significant differentially expressed features between treated vs un-treated samples within the same cell line (WT or AI), representing the effect of serum stimulation on glycopeptides and phosphopeptides. Volcano plots for **B.** WT and **C.** AI comparing the specific regulated phosphorylation sites due to serum stimulation with significance threshold at ±log2(0.58) FC and FDR > 0.05; points represent IDs of phosphorylation sites (e.g. GENE_S123), where S, T, Y corresponds to phosphorylation at serine, threonine, and tyrosine residues. **D.** Two-axis comparison of serum stimulation responses in cell lines using the same differential expression data in A. plotted on log10 FC scales; points are categorized as phosphorylation sites only significant in WT (WT*), AI (AI*), both (Both) or non-significant (NS). **E.** Pathway ORA by g:Profiler for stimulation effect on phosphorylated genes using KEGG and Reactome datasets. Genes of significant differentially expressed phosphopeptides were enriched separately for up-and down-regulation, and missing pathways were not significant. **F.** Pathway-focused heatmap built from all phosphosites of a curated gene set from the enriched pathways and cell line in E. (any gene may contain both up- and down-regulated phosphosites).

Volcano plots highlight the magnitude and direction of regulation in each cell line (**Fig. 3B and Fig. 3C**). In WT, hundreds of phosphosites pass significance thresholds, including well-known signaling nodes such as RPS6KA5 that report on mTOR/S6K axes. AI exhibits fewer significant phosphosites overall and a distinct set of regulated sites. The CEP97 protein responsible for centriolar and centrosomal organization exclusively shows a greater than 3-fold up-regulation compared to other differentially expressed phosphosites. As such, although the overall phosphorylation signaling is blunted in AI cells, the data suggest distinct regulation of phosphosites compared to WT, with potential rewiring of signaling flux rather than a simple attenuation of pathway activity. To directly compare cell-type specificity in signaling regulation, we projected the log2 FC for serum stimulated phosphosites on a two-axis scatter in **Fig. 3D**. Interestingly, we found that a substantial amount of phosphosites was only specific to WT (WT*) or AI (AI*) cell lines. The observed 444 WT specific phosphosites and 184 AI specific phosphosites may suggest that many nodes engage in a cell line or glycan-dependent manner. When we collapsed these significantly regulated phosphosites to their corresponding proteins, we also observed that there were 282 WT specific and 99 AI specific proteins in the same manner, indicating that many of the nuances are beyond variations of phosphosites on the same protein and instead reflect different protein engagements. The above data for differential analysis of serum stimulated phosphosites can be found in **Supplementary Table 3**.

To investigate the biological processes that are preferentially engaged in each cell type upon serum stimulation, we performed pathway over-representation analysis (ORA) in g:Profiler on the significant genes which passes the 1.5 FC threshold in WT and AI serum stimulation (**Fig. 3E**)^86^. We used Reactome and KEGG databases for their comprehensive coverage of mammalian signaling pathways and their regular curation with experimental validation^87,88^. Up- and down-regulated gene lists were analyzed separately, and we displayed the top 20 pathways per direction considering both cell lines after filtering to enrichment FDR < 0.05 and a non-zero intersection size. The enrichment results for all pathways can be found in **Supplementary Table 3**. In general, AI-specific genes were enriched in many fewer pathways compared to WT in both up- and down-regulated categories. AI cells exhibited notable absence in pathways like longevity, cGMP-PKG, insulin signaling, MAPK signaling, and oxytocin signaling amongst up-regulated phosphosites. The interesting observation is that although AI cells were not up-regulated in these pathways, they were also not susceptible to the down-regulation of epidermal growth factor receptor (EGFR), mammalian target of rapamycin (mTOR), ErbB, and Rho GTPase signaling as WT cells exhibited.

To examine the specific phosphosites that cause the observed enrichment pattern, we assembled heatmaps to visualize patterns in exemplar nodes sourced from biologically important pathways (**Fig. 3F**). We selected all phosphosites of a curated gene set from the enriched pathways and cell line sources in **Fig. 3E** (any gene may contain both up- and down-regulated phosphosites), many of which are selected since they were not enriched for AI. Preprocessed intensities of these phosphosites (statistically significant in **Fig. 3A**) were z-score normalized across all stimulation samples, applied with supervised clustering by enrichment pathway source, followed by un-supervised hierarchical clustering within pathway by Euclidean distance and complete linkage. Each sample was also hierarchically clustered, and we found that the dominant difference in z-scores amongst same phosphosites were by serum stimulation conditions, rather than by cell line. This observation was expected and was opposite to the global view of protein expression and PTM patterns in (**Fig. 1B**). We can further confirm that by matching the rows for the same cell line and comparing their treatment conditions, the z-score differences are dissimilar. For example, in the up-regulated heatmap EGFR_T711 enriched for the MAPK signaling pathway demonstrates a much weaker up-regulation due to serum-stimulation in AI than WT. Additional phosphosites including ITPR3_S1846, APC_S2809, RPS6KB2_S443, S431, S460, EIF4EBP1_T75, and NUP153_543, T644, T647 also show diminished responses to serum stimulation in AI cells. In the down-regulated heatmap, MTOR_S2477, as well as PRKAA2_S409 and PRKAG3_T408 on the mTOR signaling pathway, demonstrate weaker serum-induced down regulation in AI than WT. The same diminished downregulation was observed in ERBIN_S621, S636, S648, enriched from the ERBB2 signaling pathway and on the well-known on the receptor tyrosine kinase signaling axis^89^.

### Integrated network analysis reveals functional links between the glycoproteome and phosphoproteome

To identify functional connections between glycosylation remodeling and phosphorylation signaling networks, we integrated the glycoproteomics data under the starvation condition as in **Fig. 2** and phosphoproteomics data under serum stimulation as in **Fig. 3** to generate protein-protein interaction (PPI) networks. Using Cytoscape and the STRINGapp, we paired the significantly regulated glycopeptides (AI/WT) with serum stimulated phosphosites for both WT and AI cell lines (treated/un-treated)^90,91^. The dataset used for the search can be found in **Supplementary Table 4**. After STRING functional interaction enrichment (confidence score = 0.4), we retained the connected central cluster and applied Molecular Complex Detection (MCODE) clustering through the clusterMaker app, with node score cutoff of 0.1, degree cutoff of 2, and k-core of 2; cluster numbers are labelled by STRING enrichment in **Supplementary Table 4**^92,93^. The integrated networks exhibited extensive cross-layer connectivity, comprising 798 nodes and 4808 edges for WT, and 614 nodes and 3532 edges for AI (**Fig. 4**). The maximum log2 FC was not visualized for WT glycopeptide nodes and visualized in AI nodes (as indicated by the node outline color) to prioritize the level of regulation (AI/WT) that caused divergent stimulation responses in the neighboring phosphopeptides nodes. Node size reflected the abundance of detected glycopeptides and phosphopeptides for each protein, providing information on post-translational modification density. MCODE identified a total of 14 functional modules in WT cells and 8 in AI cells (**Supplementary Fig. 4A and 4B**), and STRING functional enrichment on various database revealed distinct organizational principles in receptor tyrosine kinase signaling, cytoskeletal organization, and the cell cycle machinery.

**Figure 4.**
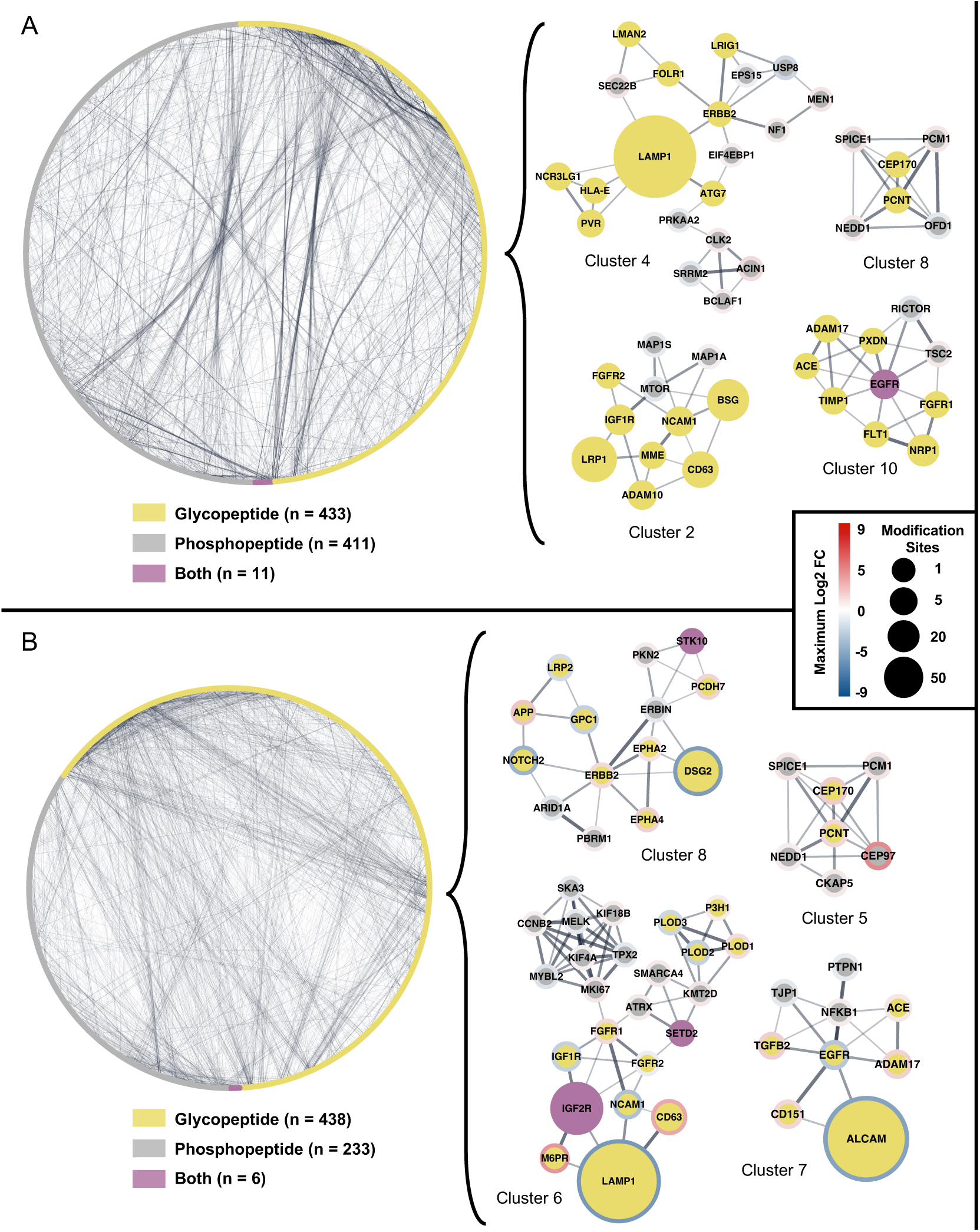
Functional interfaces between remodeled glycopeptides and serum-responsive phosphopeptides. STRING PPI of significantly regulated glycosylated proteins under starvation baseline differences (AI vs WT) combined with phosphorylated proteins under serum stimulation based on differential expression analysis. **A.** WT central cluster, and **B.** AI central cluster, arranged in circular layout, with smaller biologically relevant MCODE clusters shown. Edges in network represent proteins that are functionally connected with a minimum confidence score of 0.40, where edge thickness is proportional to confidence.

In WT cells, the network organization highlighted specific signaling and structural modules driven by key kinases and receptors (**Fig. 4A**). MCODE Cluster 10 centered on EGFR, identified as a dual glyco- and phosphopeptide (purple node), which connected to the glycosylated metalloprotease ADAM17 and the angiotensin-converting enzyme ACE (Cluster 10)^94,95^. This module also included the vascular endothelial growth factor receptor FLT1 and neuropilin NRP1, as well as mTOR regulator TSC2 and mTORC1 component RICTOR^96,97^. A separate signaling hub, Cluster 4, was organized around the phosphopeptide ERBB2; this kinase showed functional links to endocytic regulators such as USP8 and EPS15 and was anchored to LAMP1, which appeared as a large, glycosylated node (Cluster 4)^98^. The mTOR signaling machinery formed a distinct module (Cluster 2), where the kinase mTOR (phosphopeptide) connected to several surface glycoproteins, including the insulin-like growth factor receptor IGF1R, the adhesion molecule NCAM1, and the low-density lipoprotein receptor LRP1 (Cluster 2)^99^. Finally, Cluster 8 defined a centrosomal module containing the glycosylated scaffold proteins PCNT and CEP170, which were linked to the phosphorylated mitotic regulators SPICE1, PCM1, and NEDD1 (Cluster 8)^100^.

Glycoengineering in AI cells drove a significant reorganization of these functional modules and their regulatory states (**Fig. 4B**). The EGFR module (Cluster 7) shifted composition, centering on an interaction between EGFR and the activated leukocyte cell adhesion molecule ALCAM^101^. Notably, ALCAM appeared as a large, downregulated node (blue outline), while its partner ADAM17 displayed upregulation (red outline). The ERBB2 module (Cluster 8) also reorganized, linking the kinase to the desmoglein DSG2 and the amyloid precursor protein APP; DSG2 was identified as a large, strongly downregulated glycosylated node, whereas APP was upregulated (Cluster 8). The centrosome module (Cluster 5) in AI cells retained the core components PCNT and CEP170—both showing upregulation (red outline)—but recruited the centrosomal protein CEP97, which is the most strongly upregulated phosphopeptide as in **Fig. 3C**, unseen in the WT cluster^102,103^. Perhaps the most distinct feature of the AI network was the emergence of a large, multi-functional cluster (Cluster 6) that bridged receptor signaling and cell cycle machinery. This module was anchored by the upregulated dual glyco-/phosphopeptide IGF2R and the downregulated glycopeptide LAMP1^104,105^. These receptors connected to a diverse array of phosphopeptides including the fibroblast growth factor receptor FGFR1 and a sub-network of mitotic regulators such as MYBL2, MELK, and TPX2^106,106,107^.

Supplementary analysis provided further resolution on specific functional groups. Both cell lines shared a conserved extracellular matrix cluster (**Supplementary Fig. 4**, Cluster 1) enriched in laminins (LAMA2, LAMA3) and integrins (ITGA6, ITGA7)^108^. However, the WT network contained a prominent protein folding and quality control module (**Supplementary Fig. 4A**, Cluster 3) anchored by the dual glyco-/phosphopeptide HSP90B1 and the chaperone HYOU1. Conversely, the AI network uniquely exhibited a calcium signaling cluster (**Supplementary Fig. 4B**, Cluster 2) integrated by the voltage-dependent calcium channel subunits CACNA2D1, CACNA2D2, and CACNA1G, along with the inositol 1,4,5-trisphosphate receptors ITPR1 and ITPR3^109^.

## DISCUSSION

This study employed glycoengineering to perturb glycosylation pathways, phosphorylation signal stimulation, and proteomics, glycoproteomics, and phosphoproteomics frameworks to interrogate how defined alterations in N-glycan biosynthesis propagate through kinase-mediated signaling networks and reshape cellular regulatory circuitry. By engineering cells to produce predominantly non-fucosylated, highly sialylated bi-antennary glycans and coupling this genetic perturbation with serum stimulation dynamics, we provide an integrated map connecting glycan remodeling to network-wide phosphoproteome reorganization. Our multi-omics approach reveals that glycosylation acts not merely as a surface protein modification but also act as a fundamental architect of cell surface proteins to response to extracellular stimulation and intracellular signaling topology, with implications for understanding disease mechanisms and identifying therapeutic vulnerabilities in glycosylation-remodeled pathological states.

### Glycoengineering establishes system-wide molecular reorganization

Our multiplexed proteomics platform revealed that glycoengineering represents the dominant molecular signature across all three omics layers, demonstrating that targeted manipulation of glycan biosynthesis creates system-wide effects extending far beyond the glycoproteome itself. The unsupervised clustering patterns in **Fig. 1B**, with separation of AI and WT across both conditions and cell line, may suggest that glycoengineering robustly rewires the baseline molecular state across modalities, while acute serum stimulation elicits immediate and pronounced phosphorylation changes with comparatively limited short-term effects on intact glycopeptides. This pervasive influence validates our experimental design for dissecting glycan-phosphorylation regulation and underscores that cells cannot compartmentalize glycosylation changes to the secretory pathway alone. The finding that glycoengineering status shapes the molecular landscape more profoundly than acute serum stimulation has important implications for understanding how cells maintain identity despite environmental fluctuations. Glycosylation may serve as a slower acting but more persistent regulatory layer that establishes cellular state, while phosphorylation provides rapid, reversible responses within that framework. The temporal decoupling we observed between glycoproteome stability and phosphoproteome dynamics over the stimulation window supports this hierarchical model, where baseline glycosylation patterns potentially constrain the range of acute signaling responses available to cells.

### Baseline glycan remodeling establishes altered phosphorylation landscape

The baseline comparison in **Fig. 2** revealed that glycoengineering created extensive glycoproteome remodeling that successfully mimicked disease-relevant patterns of altered fucosylation, sialylation, and branching. We attribute this successful remodeling likely due to the genetic modifications on glycosyltransferase enzymes as suggested by **Fig. 2A**, where the regulation of protein expression was according to our expectations during both DIA and DDA assessments. Notably, the residual abundance detected for some knockout targets likely reflects the identification of peptides originating from truncated, loss-of-function transcripts upstream of the CRISPR-induced frameshift, rather than the presence of active enzyme. The relatively small number and magnitude of changes in phosphorylated features under baseline conditions is expected (**Fig. 2B**), as cells are un-synchronized in cell-cycle phases, basal phosphorylation may be homeostatically constrained, and many signaling nodes are quiescent in complete medium^71,72,110,111,112^. In contrast, the glycoproteome directly reflects the engineered edits at steady state, yielding the broad distribution of fold-changes we observe. Importantly, these baseline glycan changes propagated to establish substantial differences in phosphorylation state even in the absence of external stimulation, demonstrating that altered glycosylation may fundamentally reprogram the resting signaling landscape rather than simply modulate stimulus-evoked responses.

We saw that the regulation of important glycan types including fucose, sialic acid, and N-GlcNAc branching aligned with our glycoengineering efforts through assessment at the relative abundance and singular glycopeptide representation levels (**Fig. 2C, D, E and F**). Absolute percentage abundance for target glycan glasses remain moderate because a substantial fraction of MS signal derives from high-mannose or immature glycans (endoplasmic reticulum/early Golgi)^113^. As such, a residual population of fucosylated or branched structures persists even with glycoengineering because glycan maturation is asynchronous across the secretory pathway and some glycans are detected while in immature states.

By demonstrating that glycoengineering alone produces extensive and directionally coherent remodeling of glycan composition, we establish a well-defined starting state in which receptor and adhesion surfaces and ligand interactions are altered. Considering the previously summarized literature, such remodeling is expected to modulate receptor clustering, phosphodegron exposure, and kinase engagement^11,114,115,116^. The presence of baseline phosphorylation differences as indirect consequences of glycan remodeling suggests multiple possible mechanisms: altered receptor trafficking may change surface receptor density and composition, modified glycan structures may affect spontaneous receptor dimerization or clustering, altered interactions with the extracellular matrix or soluble factors may provide constitutive activation signals, or changes in glycoprotein stability may shift the balance between signaling-competent and degraded receptor pools. These mechanisms likely operate in combination, creating a new steady state where basal kinase activities are redistributed even before cells encounter growth factor stimulation.

### Glycan context dictates pathway-selective serum responses

The serum stimulation experiments demonstrated that glycoengineering not only establishes distinct baseline states but also fundamentally alters how cells interpret and transduce growth factor signals. The observation that the serum treatment effect is concentrated in the phosphoproteome, whereas the glycoproteome changed minimally over the 30-minute serum stimulation window, accords with the transient nature of phosphorylation cascades and with our baseline observation that glycan remodeling is a more permanent feature that settles in equilibrium (**Fig. 3A**). The temporal separation between these post-translational modifications reflects their distinct regulatory roles: phosphorylation operates on second-to-minute timescales enabling rapid signal transduction, while N-glycan remodeling requires much longer for complete processing due to residue accessibility and progress through the secretory pathway^117,118,119,120^. Together with the baseline findings, we may speculate that engineered glycans set the initial context of the cellular environment, and serum stimulation predominantly actuates kinase-mediated responses upon that background.

While both WT and AI cells responded to serum with widespread phosphorylation changes, the specific pathways engaged and their relative activation intensities diverged substantially. The substantially greater serum response in WT compared to AI cells, combined with the distinct set of regulated sites, suggests that glycan modifications create a comprehensive rewiring of signaling flux rather than a simple attenuation of pathway activity (**Fig. 3B and C**). CEP97 phosphorylation, which showed exclusive a more than three-fold difference in up-regulation compared to other phosphosites in AI cells, has been implicated in regulating ciliogenesis and centrosome function through kinase-dependent mechanisms, suggesting that glycan modifications may influence cell cycle checkpoint control and mitotic fidelity^103,102^.

The pathway ORA revealed that AI-specific genes were enriched in many fewer pathways compared to WT in both up- and down-regulated categories (**Fig. 3E**). AI cells exhibited notable absence in pathways like longevity, cGMP-PKG, insulin signaling, MAPK signaling, and oxytocin signaling amongst up-regulated phosphosites. This asymmetric response pattern suggests that glycan modifications may create a buffered signaling state where certain growth-promoting pathways are constitutively restrained while stress-response and homeostatic pathways remain intact. The interesting observation that although AI cells were not up-regulated in these pathways, they were also not susceptible to the downregulation of EGFR, mTOR, ErbB, and Rho GTPase signaling as WT cells exhibited may indicate selective pathway insulation rather than global signal dampening. For pathways in which both cell lines are present, AI cells demonstrate increased enrichment in SUMOylation, gene transcription and expression, mRNA transport, and chromatin modulation, as well as less decrease in enrichment of signal transduction pathways.

The examination of specific phosphosites in exemplar signaling nodes revealed that the dominant difference in z-scores amongst the same phosphosites were by serum stimulation conditions, rather than by cell line, as evident by unsupervised clustering (**Fig. 3F**). This observation was expected and opposite to the global view of protein expression and PTM patterns observed in the multi-omics overview. The clustering pattern confirms that serum stimulation produces the primary biological signal that drives phosphorylation changes, while glycan context may modulate the amplitude and duration of these responses rather than completely reorganizing the temporal sequence of pathway activation.

EGFR_T711, enriched for the MAPK signaling pathway, demonstrates much weaker up-regulation due to serum-stimulation in AI than WT. EGFR Thr phosphorylation occurs within the kinase domain and has been associated with receptor endocytosis and downstream signaling capacity^121^. Additional nodes including ITPR3_S1846, which regulates calcium release from endoplasmic reticulum stores coupling growth factor signaling to intracellular calcium dynamics, also demonstrate attenuated serum responsiveness in AI cells^122^. RPS6KB2, EIF4EBP1, and NUP214 phosphosites also show diminished responses to serum stimulation in AI cells. In the down-regulated heatmap, MTOR_S2477, as well as PRKAA2_S409 and PRKAG3_T408 on the MTORC1 signaling pathway, demonstrate weaker serum-induced downregulation in AI than WT. MTOR Ser phosphorylation has been reported to modulate mTORC1 assembly and activity in response to nutrient availability, while PRKAA2 (AMPKα2) and PRKAG3 (AMPKγ3) subunits regulate the energy-sensing capacity of AMPK complexes^97,123,124^. The attenuated downregulation in AI cells suggests that glycan modifications may stabilize MTOR pathway components against feedback inhibition, potentially through altered receptor endocytosis rates or galectin-lattice-mediated surface retention. The same diminished downregulation was observed in ERBIN_S1266 on the receptor tyrosine kinase signaling axis^89^.

These observations align with our hypothesis that glycoengineered AI cells do not merely demonstrate blunted cellular signaling, but rather a comprehensive rewiring of cellular flux. This pathway-selective rewiring indicates that glycan structures on cell surface receptors potentially influence coupling efficiency between receptor activation and downstream kinase cascades, rather than causing wholesale loss of signaling capacity.

### Integrated networks reveal glycan-dependent reorganization of signaling modules

The integrated network analysis revealed extensive reorganization of protein interaction modules in glycoengineered cells, with a reduction from 14 functional modules in WT to 8 in AI. These topological changes indicate that glycosylation establishes organizing principles determining which proteins functionally cluster to form signaling modules. In WT cells, signaling architecture followed canonical compartmentalization, with distinct modules dedicated to EGFR (Cluster 10) and mTOR (Cluster 2) signaling axes. However, glycoengineering triggered a rewiring of this signaling insulation. The collapse of discrete receptor clusters suggests that the specific glycan features lost in the AI line—namely core fucosylation and complex branching—may be essential for maintaining the spatial segregation required for independent receptor tyrosine kinase signaling^66,69^. The hyper sialylation of cell surface glycoproteins may further be associated with various ligand mediated signal which lead to downstream AKT phosphorylation and apoptosis pathways^99^.

This potential erosion of surface stability is illustrated by the remodeling of the EGFR module. In WT cells (**Fig. 4A**), EGFR exists as a central signaling hub; in AI cells (**Fig. 4B**), it reorganizes into a complex dominated by the adhesion molecule ALCAM and the metalloprotease ADAM17. The physical clustering of downregulated EGFR with upregulated ADAM17 in the glycoengineered network implies a functional coupling where altered receptor glycosylation may promote proteolytic processing or destabilization rather than canonical signaling^101^. Consequently, the network topology suggests that glycoengineered cells may shift reliance from high-fidelity growth factor sensing toward alternative adhesion-dependent survival mechanisms. Furthermore, the emergence of a multi-functional cluster (Cluster 6) in AI cells represents a significant rewiring of cellular logic connecting the cell surface to the nucleus. This module physically bridges the dual glyco-phospho hub IGF2R and the lysosomal protein LAMP1 directly with mitotic drivers (CCNB2, MELK) and chromatin modifiers (SETD2). The integration of these diverse functional classes into a single module suggests that the hyper-sialylated state of these receptors in AI cells may alter their canonical trafficking or lysosomal functions, biasing them instead toward non-canonical scaffolding roles^104,105^. By functionally coupling surface glycoprotein dynamics directly to cell cycle machinery, the glycoengineered cells appear to bypass intermediate nutrient-sensing cascades (such as the lost mTOR module), establishing a more direct link between the extracellular glycan environment and proliferative control.

Finally, the supplementary analysis illuminates a distinct shift in homeostatic priorities beyond the core signaling hubs (**Supplementary Fig. 4**). While the extracellular matrix interface—comprising laminins and integrins—remains structurally conserved across both networks, the internal regulatory machinery diverges significantly. The WT network maintains a canonical quality control module anchored by chaperones HSP90B1 and HYOU1, reflecting standard ER-associated protein folding requirements^125^. In contrast, the AI network replaces this folding axis with a dedicated calcium signaling cluster composed of voltage-dependent calcium channel subunits (CACNA2D1/2D2/1G) and IP3 receptors (ITPR1/3). This topological switch suggests that the glycoengineered state, potentially influenced by the increased negative charge of hyper-sialylation or altered membrane dynamics, moves away from chaperone-mediated buffering. Instead, these cells appear to engage calcium-dependent signaling modalities to coordinate intracellular responses and manage the structural demands of a re-engineered glycome^109,122^. Collectively, these network shifts propose that the “glycan code” potentially functions as an insulator for signaling pathways; when this code is rewritten, the proteome undergoes fundamental re-networking to sustain cellular function through alternative, adhesion- and chromatin-coupled circuits.

### Implications for disease biology and therapeutic strategies

Our findings have significant implications for understanding disease contexts where N-glycan remodeling is prevalent, particularly cancer. The tumor-associated alterations in fucosylation, sialylation, and branching that we modeled experimentally are known to correlate with malignant transformation, metastatic potential, and therapeutic resistance^64,65,68,77,105,115^. Our demonstration that such glycan changes systematically redistribute kinase activity and pathway engagement suggests that glycosylation remodeling may serve as a mechanism for cancer cells to escape normal growth controls while maintaining essential cellular functions. The specific disruption of EGFR-mTOR signaling axis we observed, coupled with the preferential engagement of chromatin and RNA regulatory pathways in glycoengineered cells, nominates these redirected signaling programs as potential therapeutic vulnerabilities in glycosylation-altered cancers^22,97,123,126^. If tumor cells with aberrant glycosylation become dependent on alternative signaling architectures to maintain proliferation, then targeting the rewired pathways rather than canonical growth factor receptors may achieve greater therapeutic efficacy. This framework could explain why some cancers with EGFR overexpression show limited response to EGFR inhibitors if concurrent glycosylation changes have already redirected signaling through alternative routes.

The integration of quantitative glycoproteomics with phosphoproteomics under controlled perturbations establishes a methodology that can be extended to other disease contexts. In congenital disorders of glycosylation, this approach could map how specific glycosyltransferase deficiencies propagate to create systems-level signaling defects, potentially identifying compensatory pathways amenable to therapeutic intervention^10^. In immunology, applying this framework to glycosylation enzymes that regulate immune checkpoint molecules could reveal how glycan remodeling coordinately affects both checkpoint expression and downstream signaling in tumor and immune cells^28,127^. The demonstration that glycosylation changes create predictable patterns of signaling network reorganization raises the possibility of using glycan profiles as biomarkers to predict which signaling pathways will be engaged in individual patients, enabling more precise selection of targeted therapies based on glycosylation status.

### Study limitations

Several limitations of our study should be acknowledged. Our use of a single cell line limits the generalizability of our findings across different cellular contexts, and the specific combination of glycoengineering edits we employed represents only one possible perturbation among many. The 30-minute stimulation window, while appropriate for capturing acute phosphorylation responses, may not reflect longer-term adaptive changes in glycoprotein trafficking or cell surface composition. Most importantly, our analysis focused on the most abundant and reliably detected features, potentially missing subtle but biologically important changes in low abundance signaling molecules. The extreme nature of our glycoengineering intervention, while valuable for establishing proof-of-principle, makes it challenging to attribute specific phenotypes to individual glycan structures versus cumulative effects of multiple glycan alterations. The correlation-based network analysis we employed identifies co-regulated proteins but does not definitively establish physical interactions or causal relationships, which would require complementary approaches such as affinity purification or genetic perturbation of individual network nodes.

### Future directions

Future studies should extend this framework to disease-relevant cell types and primary tissues, while incorporating longer time courses to capture the full temporal dynamics of glycan-phosphorylation crosstalk. The development of more selective glycoengineering approaches that target individual glycan structures or specific protein classes would enable more precise dissection of causal relationships. Integration with complementary techniques such as spatial proteomics and single-cell analysis could provide additional resolution of how glycosylation-phosphorylation crosstalk varies across cellular compartments and populations. Validation of key signaling pathway rearrangements through targeted approaches including immunofluorescence microscopy, proximity labeling, and functional assays would clarify whether the network reorganization patterns we observe through proteomics correspond to changes in spatial organization and biochemical activity. Testing whether specific glycan modifications on individual proteins are necessary and sufficient for the signaling changes we observe would establish mechanistic links between glycan structures and downstream phosphorylation networks.

More broadly, our study establishes a paradigm for systematic PTM perturbation studies that could be extended to interrogate other modification crosstalk networks, including ubiquitination, SUMOylation, and acetylation. The quantitative, multi-layered omics approach we developed generates rich datasets that capture both direct biochemical changes and downstream systems-level responses, providing the comprehensive molecular phenotyping necessary for building predictive models of cellular behavior. Such datasets could serve as valuable training resources for artificial intelligence and machine learning approaches aimed at developing virtual cell models that can predict cellular responses to genetic or pharmacological perturbations. The systematic mapping of PTM crosstalk networks through controlled perturbation studies represents a crucial step toward achieving the system-level understanding of cellular regulation required for next-generation drug discovery and precision medicine.

## MATERIALS AND METHODS

### Glycoengineering

AI cells were created using a CRISPR-Cas9 nuclease platform by sequential glycoengineering. WT adherent HEK293 host cell line obtained from the American Type Culture Collection (ATCC) were maintained at 37 °C in a humidified atmosphere containing 5 % CO₂ in high-glucose DMEM supplemented with L-glutamine, sodium pyruvate, and 10 % (v/v) fetal bovine serum. Cells were passaged every 2–3 days to keep confluence below 90 %. During passages, cels were detached with TrypLE Express. For all molecular manipulations, cells were seeded 24 h in advance to reach 70–90 % confluence and transfected with Lipofectamine 3000 according to the manufacturer’s protocol.

CRISPR/Cas9 editing used the pSpCas9(BB)-2A-GFP (PX458) backbone with single-guide RNAs designed in ChopChop to target FUT8, MGAT5, MGAT4A, MGAT4B, GMDS, and GNE. For knock-out studies, cells were co-transfected with two sgRNA/Cas9 plasmids per locus (1:1 mass ratio). In knock-in experiments at the GNE locus, the Cas9/sgRNA plasmid was co-delivered with a 240-bp donor template carrying the R263L/R266Q sialuria mutation flanked by ∼110-bp homology arms (1:2 mass ratio). Seventy-two hours post-transfection, GFP-positive single cells were isolated by fluorescence-activated cell sorting and expanded. Edited clones were verified by PCR, restriction digestion, and Sanger sequencing. For overexpression of selected glycosyltransferases, codon-optimised cassettes encoding ST6Gal1-P2A-MGAT1 and B4GalT1-T2A-MGAT2 were cloned into pBudCE4.1 under EF1α or CMV promoters, respectively. Transfected cells were selected with zeocin (200 µg mL⁻¹) for stable pool, and single clones were screened by Sambucus nigra agglutinin lectin blotting to confirm enhanced α2,6-sialylation.

### Serum Stimulation Study

Serum-starvation experiments were performed with the HEK293 WT cell line and the engineered AI derivative cell line. Three culture conditions were examined in biological triplicate (separate culture dish per replicate): i) normal growth in complete medium, ii) 12 h serum starvation, and iii) 12 h serum starvation followed by 30 min re-addition of 10 % FBS.

Cryopreserved cells were revived into DMEM supplemented with 10 % FBS, GlutaMAX, and zeocin (200 µg mL⁻¹). After two to three passages under antibiotic selection, cells were expanded first in T-flasks and then in 150 mm dishes until cultures reached approximately 2 × 10⁷ cells per dish. For starvation, complete medium was replaced with serum-free DMEM containing GlutaMAX and cells were incubated overnight (≈ 12 h). For the re-feeding condition, 10 % FBS was re-introduced for 30 min immediately before harvest. Cultures were transferred to ice, washed twice with ice-cold PBS, and cells were detached by gentle scraping. Cell suspensions were pelleted at low speed, washed once more with cold PBS, and the final pellets were flash-frozen and stored at −80 °C pending downstream analyses.

### Sample processing for protein extraction and tryptic digestion

The lysis buffer was prepared with 8 M urea, 75 mM NaCl, 50 mM tris(hydroxymethyl)aminomethane (Tris) pH 8.0, 1 mM ethylenediaminetetraacetic acid (EDTA), 2 μg/ml aprotinin, 10 μg/ml leupeptin, 1 mM phenylmethanesulfonyl fluoride (PMSF), 10 mM NaF, 1:100 phosphatase inhibitor cocktails 2 and 3 (PIC2 and PIC3), and 20 μM O-(2-Acetamido-2-deoxy-D-glucopyranosylidenamino) N-phenylcarbamate (PUGNAc). Each cell pellet was kept on ice and mixed with 100 μl of chilled lysis buffer. The mixture was vortexed for 20 seconds on the highest setting and the cells were lysed at 4 °C with the chilled lysis buffer for 15 minutes. This process was repeated, resulting in a total of four lysis cycles. The lysates were then centrifuged at 4 °C (20,000 g) for 10 minutes to pellet cell debris. The supernatants, containing solubilized proteins, were transferred to new 1.7 ml conical tubes. The protein concentrations in the supernatant were estimated by BCA assay (Pierce). Diluting the samples, a water blank, and lysis buffer samples 1:20 and running them in triplicate alongside a BSA standard curve further assessed the protein concentration. After incubating the samples at 37 °C for 30 minutes, the absorbance was measured at 562 nm. The sample concentrations were adjusted to 5 μg/μl with lysis buffer, and 500 μg protein aliquots of each sample were prepared for subsequent processing. The aliquoted proteins were first reduced by incubation with a final concentration of 6 mM dithiothreitol (DTT) for 1 hour at 37 °C. The reduced proteins were subsequently alkylated with 12 mM iodoacetamide (IAA) for 45 minutes at 25 °C in the dark. To decrease the urea concentration below 2 M, each sample was diluted 1:4 with 50mM Tris HCl pH 8.0. The diluted samples were then digested by adding LysC (Wako Chemicals) at an enzyme to substrate ratio of 1:50, followed by a 2-hour incubation at 25 °C. Following the LysC digestion, sequencing-grade modified trypsin (Promega) was added at an enzyme to substrate ratio of 1:50, and the samples were then incubated overnight (approximately 14 hours) at 25 °C. After the digestion, the samples were acidified by adding 50% formic acid to achieve a final concentration of 1% formic acid, which brought the pH of the solution to 3. The samples were then diluted up to 1 ml with 0.1% formic acid (FA). This step precipitated any undigested protein, which was subsequently removed by centrifugation (20,000 g) for 15 minutes and discarding the pellet. Tryptic peptides were desalted on reversed-phase C18 SPE columns (SepPak, Waters) and dried using a Speed-Vac (Thermo Scientific).

### TMT labeling of peptides

The Tandem Mass Tag (TMT) 18 labeling of peptides was performed to label 250 μg of peptides per TMT channel^128^. To label 250 μg of peptides, an initial aliquot of the peptides, as measured by a peptide-level BCA assay, was prepared in 100 μl of 100 mM HEPES pH 8.5. After thorough vortexing and a brief centrifugation to mix, 500 μl of anhydrous acetonitrile was added to 5 mg of TMT18 reagents (Thermo Fisher Scientific). This mixture was left to sit for 5 minutes, then vortexed and centrifuged. Next, 50 μl of the prepared TMT reagent (equivalent to 500 μg) was added to each corresponding tube containing 250 μg of peptides, maintaining a peptide to TMT reagent ratio of 1:2. The reaction mixture was incubated at room temperature shaking at 1,000 rpm for 1.5 hours. Following the incubation, the reactions were quenched with 6 μl of 5% hydroxylamine at room temperature for 15 minutes with shaking. The differently labeled samples were then combined. The labeled peptides were dried down and reconstituted with 3 ml of 3% acetonitrile (ACN)/0.1% FA. The pH was assessed with indicator paper, and if necessary, adjusted to pH 3 with formic acid. Finally, the sample was cleaned up using SepPak.

### Peptide fractionation

The dried peptides were reconstituted in 900 μl of 20 mM ammonium formate (NH4HCO2, pH 10) and 2% ACN and loaded onto a 4.6 mm x 250 mm RP Zorbax 300 A Extend-C18 column with 3.5 μm size beads (Agilent). Peptides were separated at a flow-rate of 1 ml/min using an Agilent 1200 Series HPLC instrument via basic reversed-phase liquid chromatography (bHPLC) with Solvent A (2% ACN, 5 mM NH4HCO2, pH 10) and a non-linear gradient of Solvent B (90% ACN, 5 mM NH4HCO2, pH 10) as follows: 0% Solvent B (9 min), 6% Solvent B (4 min), 6% to 28.5% Solvent B (50 min), 28.% to 34% Solvent B (5.5 min), 34% to 60% Solvent B (13 min), and holding at 60% Solvent B for 8.5 min. Collected fractions were concatenated into 24 fractions by combining four fractions that are 24 fractions apart (i.e., combining fractions #1, #25, #49, and #73; #2, #26, #50, and #74; and so on); a 5% aliquot of each of the 24 fractions was used for global proteomic analysis, dried in a Speed-Vac, and resuspended in 3% ACN/0.1% FA prior to LC-MS/MS analysis. The remaining sample was utilized for phosphopeptide enrichment.

### Enrichment of phosphopeptides by Fe-IMAC

The remaining 95% of the sample was further concatenated into 12 fractions (i.e., combining fractions (#1 and #12, #2 and #13 …) before being subjected to phosphopeptide enrichment using immobilized metal affinity chromatography (IMAC) as previously described. In brief, Ni-NTA agarose beads were used to prepare Fe3+-NTA agarose beads, and 300 μg of peptides were reconstituted in 80% ACN/0.1% trifluoroacetic acid (TFA) and incubated with 10 μl of the Fe3+-IMAC beads for 30 min. Samples were then centrifuged, and the supernatant containing unbound peptides was removed. The beads were washed twice and then transferred onto equilibrated C-18 Stage Tips with 80% ACN/0.1% TFA. Tips were rinsed twice with 1% FA and eluted from the Fe3+-IMAC beads onto the C-18 Stage Tips with 70 μl of 500 mM dibasic potassium phosphate, pH 7.0 a total of three times. C-18 Stage Tips were then washed twice with 1% formic acid, followed by eluting of the phosphopeptides from the C-18 Stage Tips with 50% ACN/0.1% formic acid twice. Samples were dried down and resuspended in 3% ACN/0.1% FA prior to LC-MS/MS analysis.

### Enrichment of glycopeptides by MAX enrichment

IMAC flow-through fractions were acidified in 0.1 % formic acid (FA, pH < 3) and desalted on Sep-Pak C18 cartridges previously conditioned with 100 % acetonitrile (ACN), 50 % ACN/0.1 % FA and 0.1 % trifluoroacetic acid (TFA). After the samples were loaded twice and washed with 0.1 % TFA, peptides were sequentially eluted with 50 % and 95 % ACN/0.1 % TFA. The combined eluate was brought to ∼95 % ACN/1 % TFA and applied to Oasis MAX mixed-mode anion-exchange cartridges^47^. MAX cartridges were equilibrated with ACN, 100 mM triethylammonium acetate, water, and 95 % ACN/1 % TFA before the sample was loaded twice. Non-glycosylated peptides were removed by washing with 95 % ACN/1 % TFA, and intact glycopeptides were eluted with 50 % ACN/0.1 % TFA. Enriched fractions were dried, reconstituted in 3 % ACN/0.1 % FA, and prior to LC-MS/MS analysis.

### LC-MS/MS analysis

Samples were analyzed using either an Orbitrap Ascend coupled to an EvoSep One (DDA mode) or a timsTOF HT coupled to an EvoSep One (DIA mode). Chromatographic separation was consistent across platforms, utilizing a PepSep C18 column (15 cm x 150 µm, 1.5 µm, Bruker) heated to 50 °C within a Bruker column toaster, operating under a 30 SPD gradient.

For TMT-labeled global and phosphoproteomic fractions, the Orbitrap Ascend operated with a 2 s cycle time. MS1 scans (60,000 resolution, 350–1800 m/z, AGC 4.0e5) were followed by MS2 acquisition at 50,000 resolution. Precursors were isolated with a 0.7 m/z window and fragmented via HCD (energy 34%). Glycoproteomic fractions were analyzed using distinct parameters: MS1 range was 500–2000 m/z, and MS2 fragmentation utilized stepped HCD (25, 35, 45%) with scans acquired at 30,000 resolution to optimize glycopeptide detection.

Global DIA analysis on the timsTOF HT utilized the diaPASEF method. The scan range covered 100–1700 m/z (MS1) and 338–1338 m/z (MS2), with an ion mobility range (1/K0) of 0.70–1.45 V·s/cm² and a ramp time of 85.0 ms.

### Identification and quantification of phosphopeptides, glycopeptides, and proteins

DDA datasets were searched against the human UniProt database using MS-PyCloud (global/phospho) or GPQuest (glyco)^129^. Searches assumed trypsin digestion (max 2 missed cleavages), with fixed Cys-carbamidomethylation and TMT-labeling, and variable Met-oxidation. Phosphorylation (S, T, Y) was considered variable for phosphoproteomic searches. A target-decoy approach ensured a <1% FDR. TMT reporter ion intensities were corrected using manufacturer-supplied factors.

DIA datasets were analyzed via Spectronaut 19.1. The search archive generation matched the DDA enzymatic constraints (length 7–52 AA). Variable modifications included Met-oxidation and N-terminal acetylation, with an N-to-D deamidation modification added for glycosite library generation. Identification and quantification were performed without cross-run normalization, maintaining a 1% FDR at the PSM, peptide, and protein levels.

### Bioinformatics and Data Analysis

All data analysis and figure generation were performed using R (version 4.3.1) unless otherwise specified. BioRender was used for schematic diagrams. GraphPad Prism was used for selected bar plots. Cytoscape (version 3.10.1) along with STRINGapp was used for network interaction and visualization^90,91^. Raw proteomics data underwent systematic preprocessing to ensure analytical robustness and cross-sample comparability. All datasets (global proteome, glycoproteome, and phosphoproteome) were log2-transformed and subjected to robust median-centering normalization using PhosR-like functions. Missing values were imputed using ensemble imputation via DreamAI.

Differential expression analysis for glycoengineering efficacy on global DIA and DDA datasets was done by a one-tailed student’s t-test, with a significance threshold of p < 0.05. Differential expression analysis for all other analyses was performed using an empirical-Bayes linear-model framework implemented in the Limma package. For pairwise comparisons (AI vs WT under control conditions), moderated t-statistics were computed with variance shrinkage across features. For multi-group comparisons, one-way ANOVA was employed to identify features varying across the full experimental design (cell line × condition). Statistical significance was defined as false discovery rate (FDR) < 0.05 using the Benjamini-Hochberg procedure, combined with an absolute FC threshold of > 0.58 (corresponding to ≈1.5-fold change). For cross-dataset integration, the top 500 features per dataset were selected based on overall F-statistic ranking, yielding up to 1,500 features prior to completeness filtering (≥70% observed values required). These features were z-score normalized followed by concatenation into data blocks (Global, Glyco, Phospho).

Functional enrichment analysis was conducted using over-representation analysis (ORA) on significantly regulated gene sets. Pathway databases included Kyoto Encyclopedia of Genes and Genomes (KEGG), and Reactome. Up- and down-regulated gene lists were analyzed separately using hypergeometric testing with multiple testing correction via the Benjamini-Hochberg method. Enriched pathways were filtered to adjusted p-value < 0.05 with non-zero gene set intersection. The top 20 pathways per direction considering both cell lines after filtering to enrichment FDR < 0.05 and a non-zero intersection size were displayed. If one cell line has a pathway in the top 20, the other cell line is automatically considered for comparison, and if the pathway is missing it was not significant. Results were visualized as dot plots encoding statistical significance (-log10 adjusted p-value) and gene set size. Physical protein-protein interaction networks were constructed using the STRING database (version 11.5) with a confidence score threshold of 0.4. Significant glycopeptides (control condition) and phosphosites (serum stimulation) were collapsed to unique gene symbols and mapped to STRING interactions. The largest connected component was retained for each cell line and subjected to Molecular Complex Detection (MCODE) analysis using the clusterMaker2 Cytoscape plugin with node score cutoff of 0.1, degree cutoff of 2, and k-core of 2. Heatmaps were generated using hierarchical clustering with Euclidean distance metrics and complete linkage. Volcano plots displayed log2 FC vs -log10 p-value with significance thresholds indicated by dashed reference lines. Statistical analyses and visualizations were implemented using ggplot2, pHeatmap, and custom R functions.

## DATA AVAILABILITY

The mass spectrometry proteomics data have been deposited to the ProteomeXchange Consortium via the PRIDE45 partner repository with the data set identifier PXD071742. Reviewers can access the dataset by logging in to the PRIDE website using the following account details:

Username

Password: fOUvgH429IKM

## CONTRIBUTIONS

E.W. conducted the majority of the experiments, analyzed the data, and wrote the manuscript. H.L. contributed to the proteomics experiments and interpreted the results. M.J.B. contributed to the study design and supervised cell line generation. H.Z. conceived and supervised the study, secured funding. All authors discussed the results and commented on the manuscript.

## ACKNOWLEDGMENTS

This work was supported by National Institutes of Health, National Cancer Institute, the Clinical Proteomic Tumor Analysis Consortium (CPTAC, U24CA271079), the Early Detection Research Network (EDRN, U2CCA271895), Pancreatic Cancer Detection Consortium (PCDC, U01CA274514). This manuscript is the result of funding in whole or in part by the National Institutes of Health (NIH). It is subject to the NIH Public Access Policy. Through acceptance of this federal funding, NIH has been given a right to make this manuscript publicly available in PubMed Central upon the Official Date of Publication, as defined by NIH.

---

**Supplementary Figure 1.**
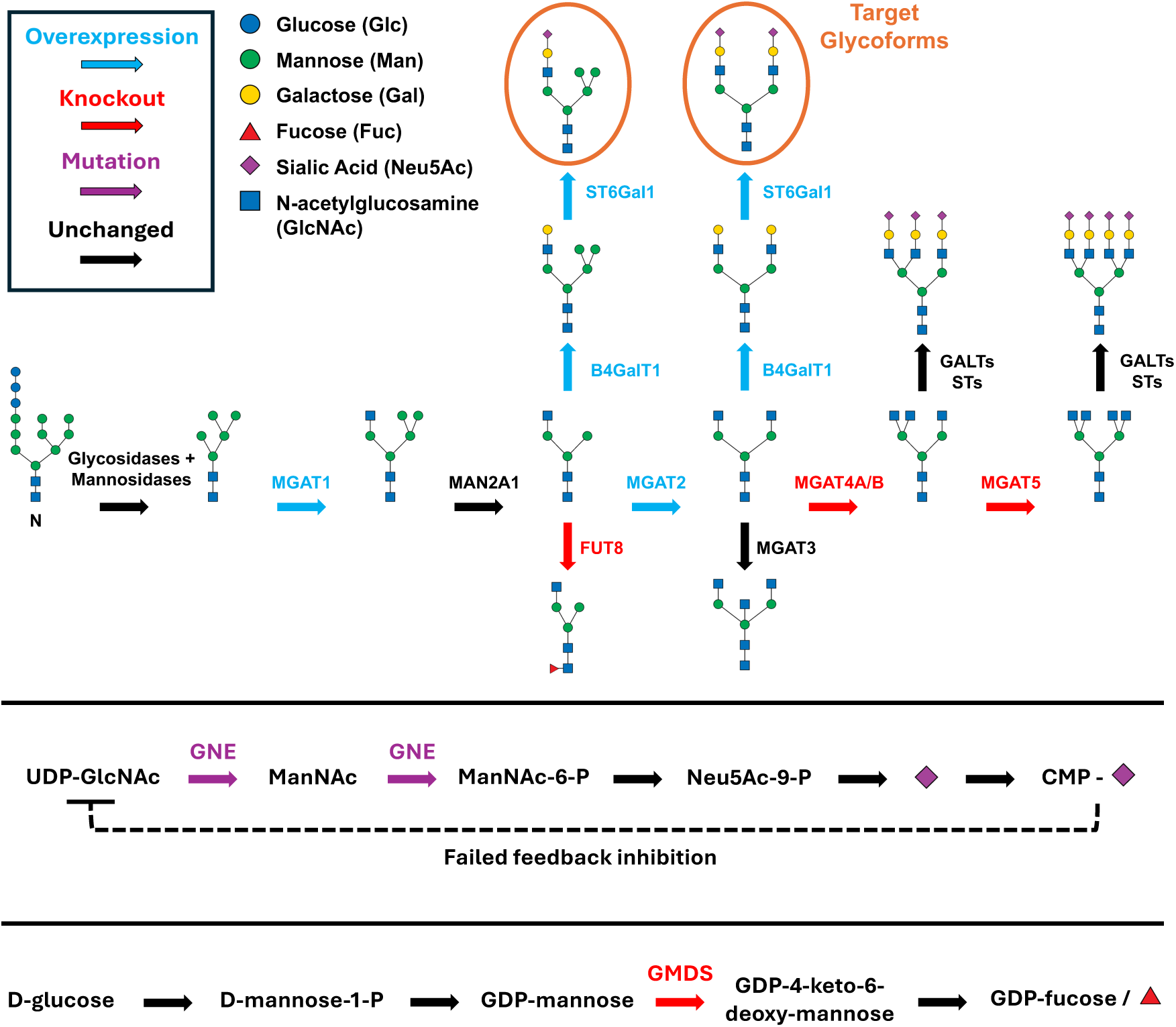
Glycoengineering Schematic of this study in the ER-Golgi landscape. Genetic modifications are shown with colored arrows including overexpression, knockout, and mutation on the corresponding enzyme that catalyzes sugar addition; GALTs represent galatosyltransferases and STs represent sialyltransferases; circled glycoforms represent the engineered dominant target hybrid and target complex glycans. The GNE mutation for sialuria and GMDS knock-out schematic is also shown.

**Supplementary Figure 2.**
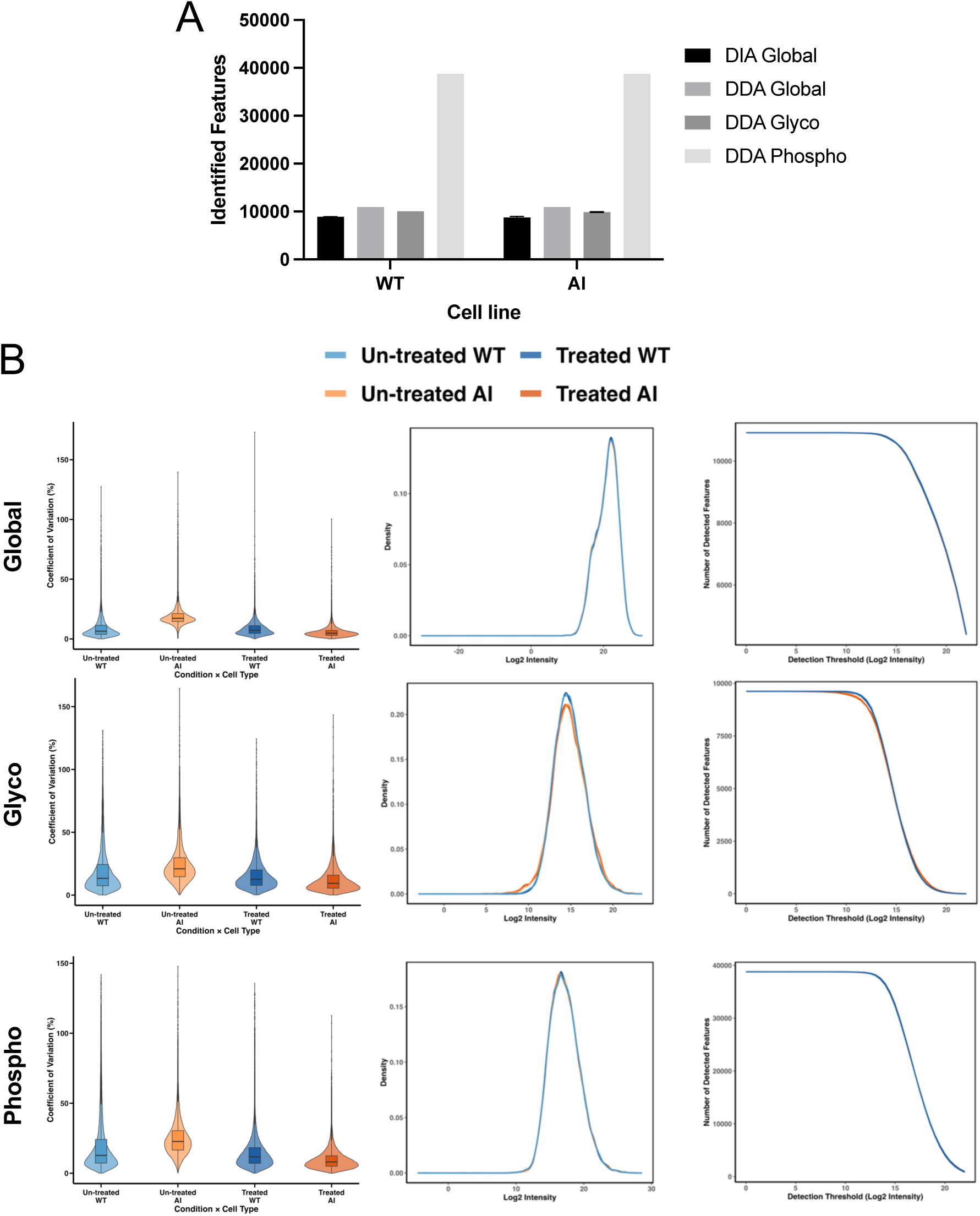
**A.** Total identified features by reads in LC-MS/MS across both cell lines (WT, AI) and conditions (un-treated, treated) in DIA global, DDA global, glyco-, and phospho-datasets; some error bars are too small to be visualized due to the relative scale. **B.** Quality control for DDA global, glyco- and phospho-datasets including CV analysis, Gaussian kernel density estimation, and feature detection thresholds.

**Supplementary Figure 3.**
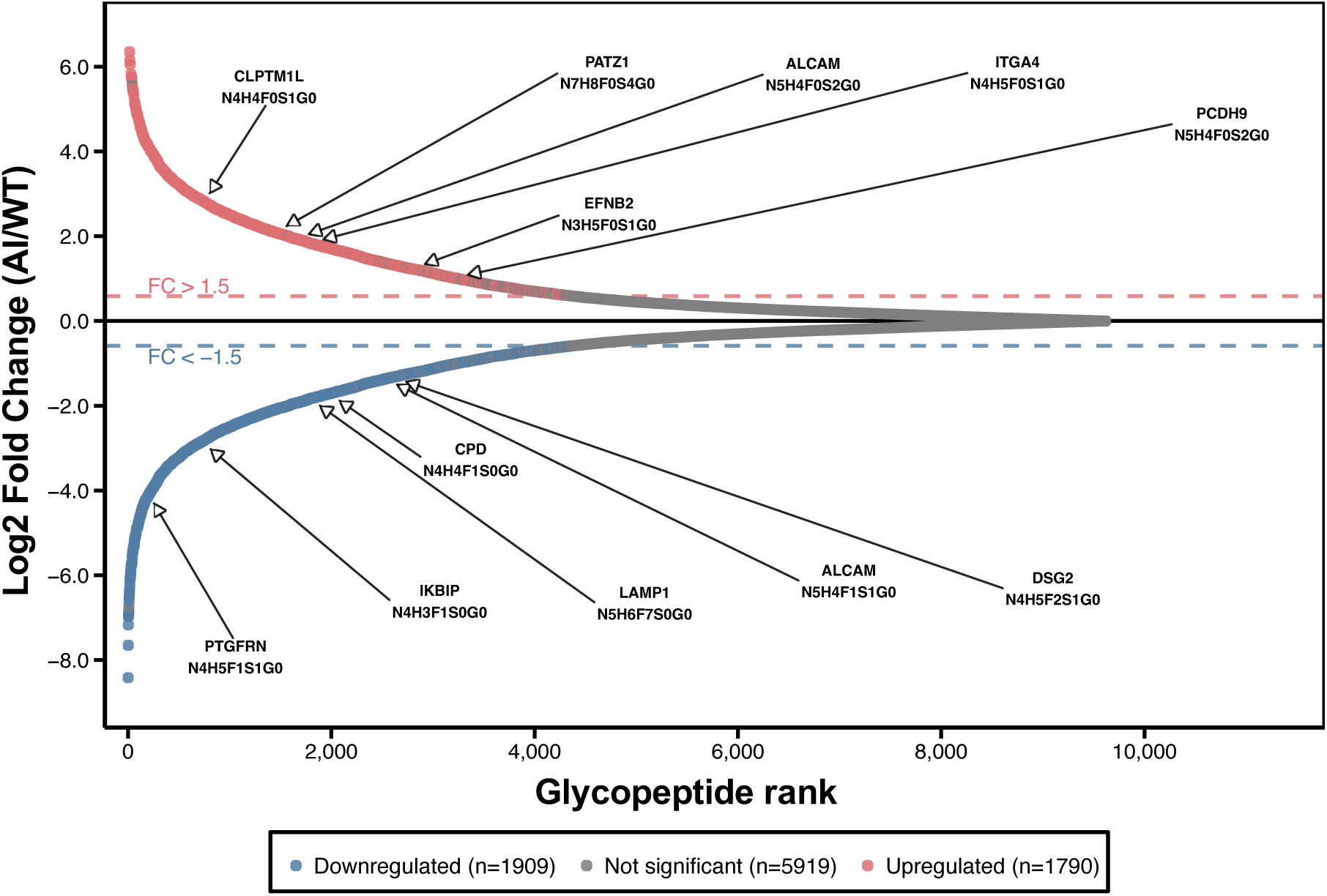
S-plot of site-resolved glycopeptide analysis depicting regulation of various fucosylated, non-fucosylated, α2,6-sialylated, and non-sialylated glycopeptides labelled by corresponding gene.

**Supplementary Figure 4.**
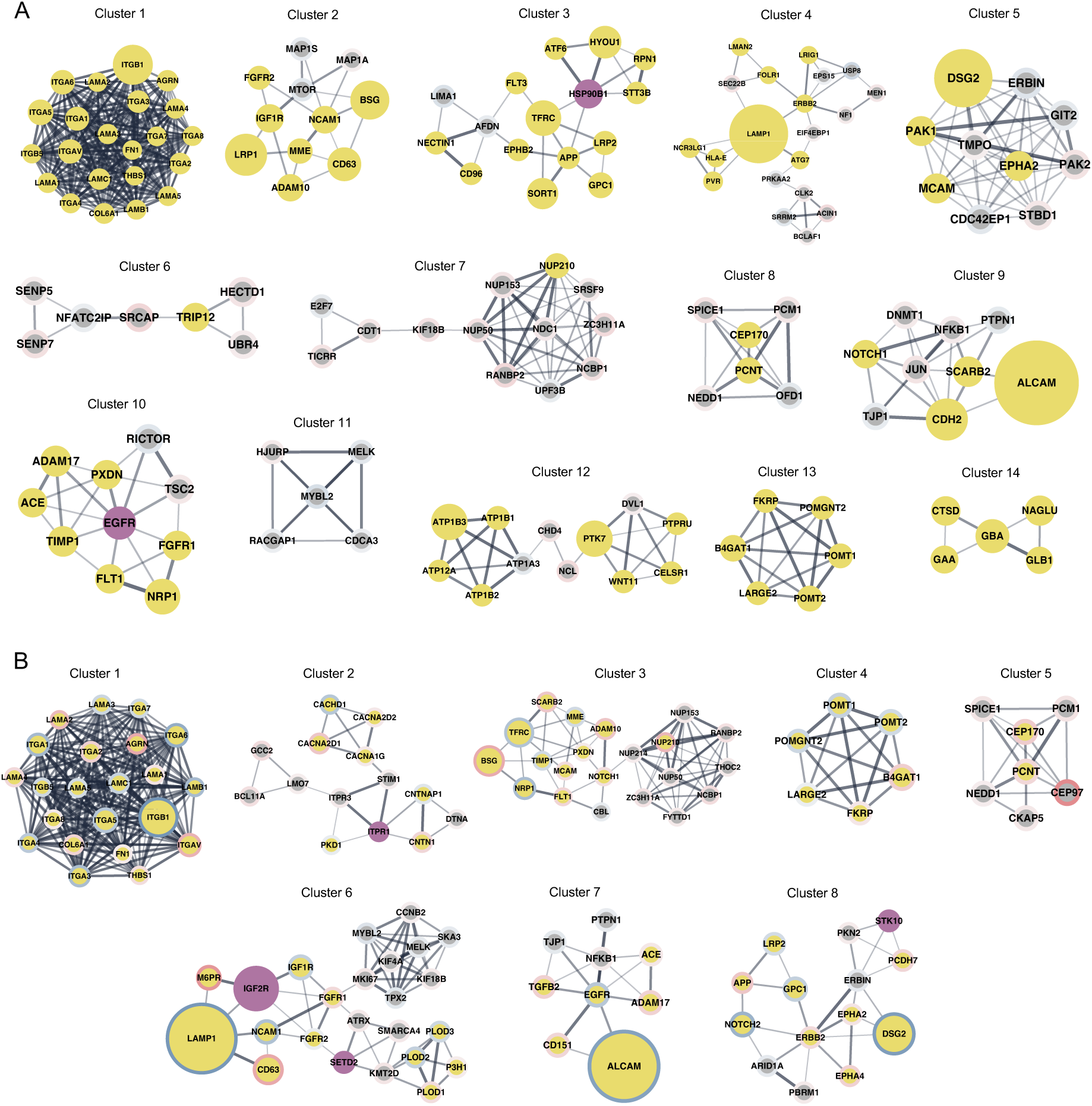
All **A.** WT and **B.** AI MCODE cluster networks identified in Cytoscape. STRING Protein-Protein Interaction (PPI) extension of main Fig. 4 with similar legend correspondence, with the exception that node sizes are not to scale as in the main figure.

